# A preclinical randomized multicenter trial of anti-IL-17A treatment for acute ischemic stroke

**DOI:** 10.1101/2021.09.29.462381

**Authors:** Mathias Gelderblom, Simon Koch, Jan-Kolja Strecker, Carina Jørgensen, Lidia Garcia-Bonilla, Peter Ludewig, Christian Bernreuther, Hans Pinnschmidt, Thiruma V. Arumugam, Goetz Thomalla, Cornelius Faber, Jan Sedlacik, Christian Gerloff, Jens Minnerup, Bettina H. Clausen, Josef Anrather, Tim Magnus

**Affiliations:** Department of Neurology, University Medical Center Hamburg-Eppendorf; Hamburg, Germany; Department of Neurology with Institute of Translational Neurology, University of Münster; Münster, Germany; Department of Neurobiology Research, Institute of Molecular Medicine, University of Southern Denmark; Odense, Denmark; Feil Family Brain and Mind Research Institute, Weill Cornell Medicine, New York; New York, USA; Department of Neuropathology, University Medical Center Hamburg-Eppendorf; Hamburg, Germany; Institute of Medical Biometry and Epidemiology, University Medical Center Hamburg-Eppendorf; Hamburg, Germany; Department of Physiology, Anatomy & Microbiology School of Life Sciences, La Trobe University; Melbourne, Australia; Translational Research Imaging Center, Clinic of Radiology, University of Münster; Münster, Germany; Biomedical Engineering Department, Centre for the Developing Brain, School of Biomedical Engineering & Imaging Sciences, King’s College London; London, United Kingdom

## Abstract

Multiple consensus statements have called for preclinical randomized controlled trials (pRCT) to improve translation in stroke research. Here, we investigated the efficacy of IL-17A neutralizing antibodies in a multicentric pRCT using a murine stroke model. C57/Bl.6 mice were subjected to transient middle cerebral artery occlusion (tMCAO). Mice were randomly allocated (1:1). Either anti-IL-17A (500 µg) or isotype antibody (500 µg) were administered 1 h after tMCAO. Primary analysis of infarct volumes was done by MRI after three days. Secondary analysis included mortality, neurological score, neutrophil infiltration and the impact of the gut microbiome on treatment effects. Out of 136 mice, 109 mice were included in the analysis. Mixed model analysis revealed that the IL-17A neutralization significantly reduced infarct sizes (anti IL-17A: 61.77 ± 31.04 mm^3^; IgG control: 75.66 ± 34.79 mm^3^; p=0.01). Secondary outcome measures showed a decrease in mortality (Hazard Ratio=3.43, 95% CI = 1.157 - 10.18; p=0.04) and neutrophil invasion into ischemic cortices. There was no difference in the neurological score. The analysis of the gut microbiome showed significant differences between centers. Taken together, this is the first positive pRCT in an ischemia reperfusion model. It suggests IL-17A neutralization as a potential target in stroke.

## Introduction

The burden of stroke on society is substantial. Stroke is the leading cause of long term disability in the Western World and an important reason for death worldwide (Collaborators, 2019). Currently, therapy is limited to thrombolysis and thrombectomy, which is only available in specialized centers. These facts clearly underline the need for novel therapeutic approaches. However, it has been challenging to translate experimental findings from preclinical observations to clinical application (Baker, 2016; O’Collins et al., 2006; Schmidt-Pogoda et al., 2020). These translational difficulties have resulted in 2010 in the first ARRIVE guidelines and a recently updated version ARRIVE 2.0 (Percie du Sert et al., 2020). A broad range of scientists have supported and extended these guidelines with calls for rigorous study designs including preclinical trials (Dirnagl et al., 2013; Landis et al., 2012). Conditions for multicenter preclinical randomized controlled trials (pRCT) are: i) harmonized experimental protocol, ii) sample size calculation, iii) randomization, iv) blinding, v) cross-validation, vi) centralized study organization. With these prerequisites, such pRCTs are hard to organize and cost intensive. Therefore, only one such trial exists in experimental stroke research (Llovera et al., 2015). Recent immunological research has indicated that neuroinflammatory responses are promising therapeutic targets in stroke (Chamorro, Dirnagl, Urra, & Planas, 2016). One of these targets is the highly conserved pro-inflammatory cytokine Interleukin-17A (IL-17). IL-17A is implicated in the pathology of several autoimmune diseases such as rheumatoid arthritis, and psoriasis, which led to FDA approved anti-IL-17 treatments (Cua & Tato, 2010; Veldhoen, 2017). In experimental stroke, evidence for a detrimental role of IL-17A was seen in multiple single-center studies (Benakis et al., 2016; Dai et al., 2020; Gelderblom et al., 2012; Lee et al., 2020; Shichita et al., 2009). While these data support an important role of IL-17A in stroke, single center studies are biased by site-specific confounders, including animal housing conditions, the microbiome, the experimental setup, and even the investigators.

Therefore, we have executed the first pRCT for IL-17A neutralization in experimental stroke. This study was carried out in Odense (Denmark), New York (USA), Hamburg (Germany) and Münster (Germany) using the temporal middle cerebral artery occlusion model (tMCAO). Following a predefined, randomized, and blinded protocol, application of IL-17A antibodies significantly reduced infarct size and mortality. Anti-IL-17A treatment resulted in less cortical neutrophil infiltration. Sequencing the gut microbiome of mice from the different participating laboratories revealed substantial differences of segmented filamentous bacteria (SFB), which are potent inducers of IL-17A producing T cells. This pRCT showed that neuralization of IL-17A is effective in a large sample size and at different study centers.

## Results

### Study design

A total of 136 C57Bl.6 mice were included in four independent research centers in Odense (Denmark), New York (USA), Hamburg, and Münster (Germany) (Figure 1). tMCAO was conducted for 45 minutes using the intraluminal filament method (6-0 nylon) (Gelderblom et al., 2012). Coded antibodies (500 μg of anti-IL-17A (Clone MM17F3, eBioscience) or 500 μg of IgG control) and filaments were provided by Hamburg. All of the participating surgeons and animal caretakers were blinded for the treatment groups. The exclusion criteria were predefined in the study protocol (Supplemental figure 1). Mice were allocated to the groups by the coordinating center using a randomizer tool. All researchers were blinded with respect to the treatment groups. Mice were randomized to treatment with either anti-IL-17A or isotype IgG antibody, which were injected intravenously 1 h after occlusion. Primary endpoint was infarct size on day three in post-mortal assessed MRI scans, secondary endpoints included mortality, and neurological scores. Only mice with a reduction of the ipsilateral blood flow below 20% of the contralateral side as measured by laser Doppler were included. Body temperature was controlled during surgery and kept at 36°C. Three days following tMCAO, mice were sacrificed, brains were removed, and sent to the core laboratory in Hamburg for evaluation of infarct size by two independent blinded investigators. Unblinding was performed following completion of the statistical analyses. In the anti-IL-17A group, eight mice were excluded for the following reasons: major intracerebral bleeding (n=2), undetectable infarct on the MRI (n=3) or technical causes (n=3). In the isotype IgG control group, six mice were excluded for the following reasons: major intracerebral bleeding (n=2), undetectable infarct on the MRI (n=1) or technical causes (n=3). The mortality until day 3 was n=3 in the IL-17A treatment group and n=10 in the isotype control group. A total of 109 mice was included in the analysis of the primary endpoint (IL-17A treatment group n=57; isotype control group n=52).

**Figure 1.**
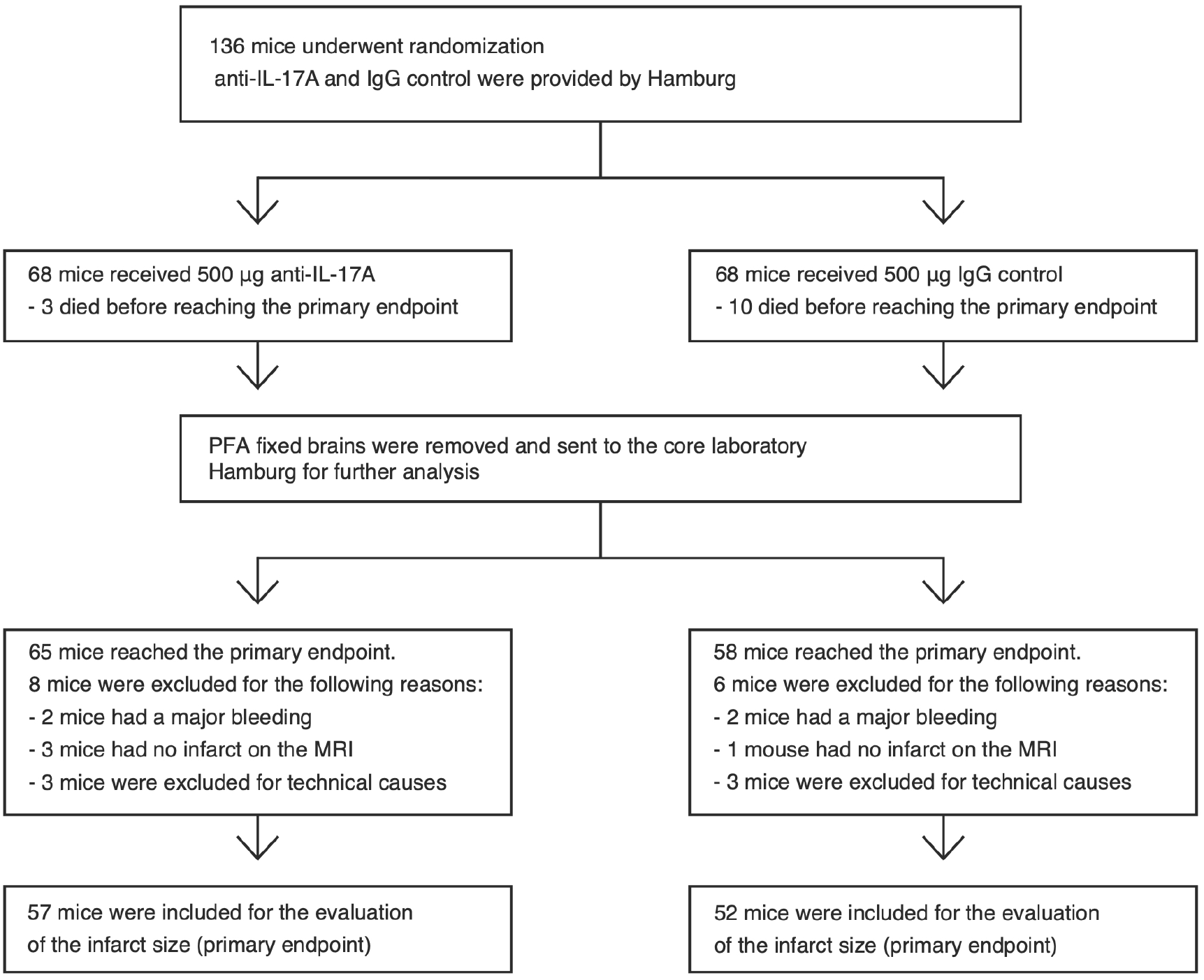
Design of the preclinical randomized multicenter study. Illustration of included and excluded animals. Mice were treated either with 500 μg of anti-IL-17A or 500 μg of IgG control 1 h following reperfusion. Randomization of antibodies and blinded analysis of infarct volumes were performed in the core laboratory in Hamburg. Only brains of mice that survived until day three underwent final analysis of infarct volumes.

### Neutralization of Interleukin-17A protects from ischemic stroke and reduces mortality

Infarct sizes were determined in a blinded approach by the volume of the apparent diffusion coefficient (ADC) lesion in post mortem brains as assessed by magnetic resonance imaging (MRI) scans. Three days following tMCAO, the mean infarct volume and SD in the pooled data set was 61.77 ± 31.04 mm^3^ in the anti-IL-17A treatment group and 75.66 ± 34.79 mm^3^ in the IgG control group. To analyze the effects of the IL-17A neutralization, we employed a linear-mixed model analysis with the two investigators as random effects, which revealed significantly reduced infarct sizes in anti-IL-17A treatment group when compared to the IgG control group (Figure 2A, B). Furthermore, mortality was significantly reduced in the anti-IL-17A group (Figure 2C, D). The probability of mice of the IgG control group to die was more than three times higher than in the anti-IL-17A group (Hazard Ratio=3.43, 95% CI = 1.157 - 10.18). Neurological scores (Supplemental Figure 2A) and weight loss until day three (Supplemental Figure 2B) did not differ between groups.

**Figure 2.**
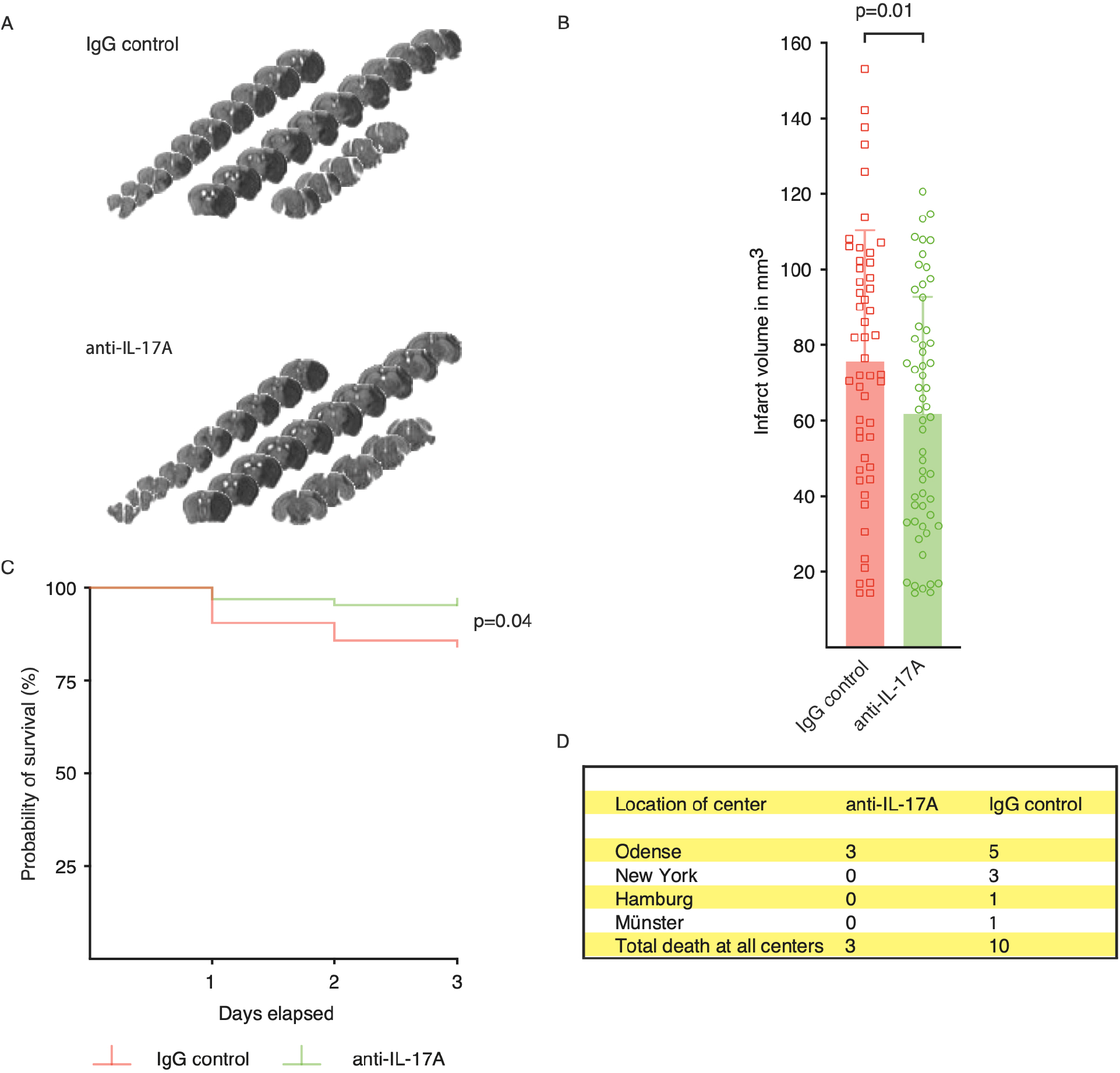
Neutralization of IL-17A significantly reduced infarct size and lowered mortality. (A) Representative MRI images of ADC weighted images. (B) Pooled data from all test centers. Infarct volumes were analyzed three days following tMCAO by MRI. The infarct volumes of rater one are shown in mm^3^. Graphs show mean±SDs. Statistical significance was assessed by linear-mixed model analysis (F_(1,209)_=6.56, p=0.01, n=109). (C, D) Survival rate and number of dead mice. Statistical significance was assessed by a log-rank test (*χ*^2^_(1)_=4.15, p=0.04, n=123).

### IL-17A neutralization reduced lesion sizes of cortical infarcts

Next, we analyzed the effects of the IL-17A neutralization on infarcts with or without involvement of cortical regions. We distinguished between lesions that affected the cortex and striatum and lesions that were limited to striatal areas and were therefore smaller on average. The pooled data showed that the neutraxlization of IL-17A significantly reduced infarcts with cortical involvement (Figure 3A). In contrast, an analysis of purely striatal infarcts showed no trend towards smaller infarcts in the anti-IL-17A group (Figure 3B).

**Figure 3.**
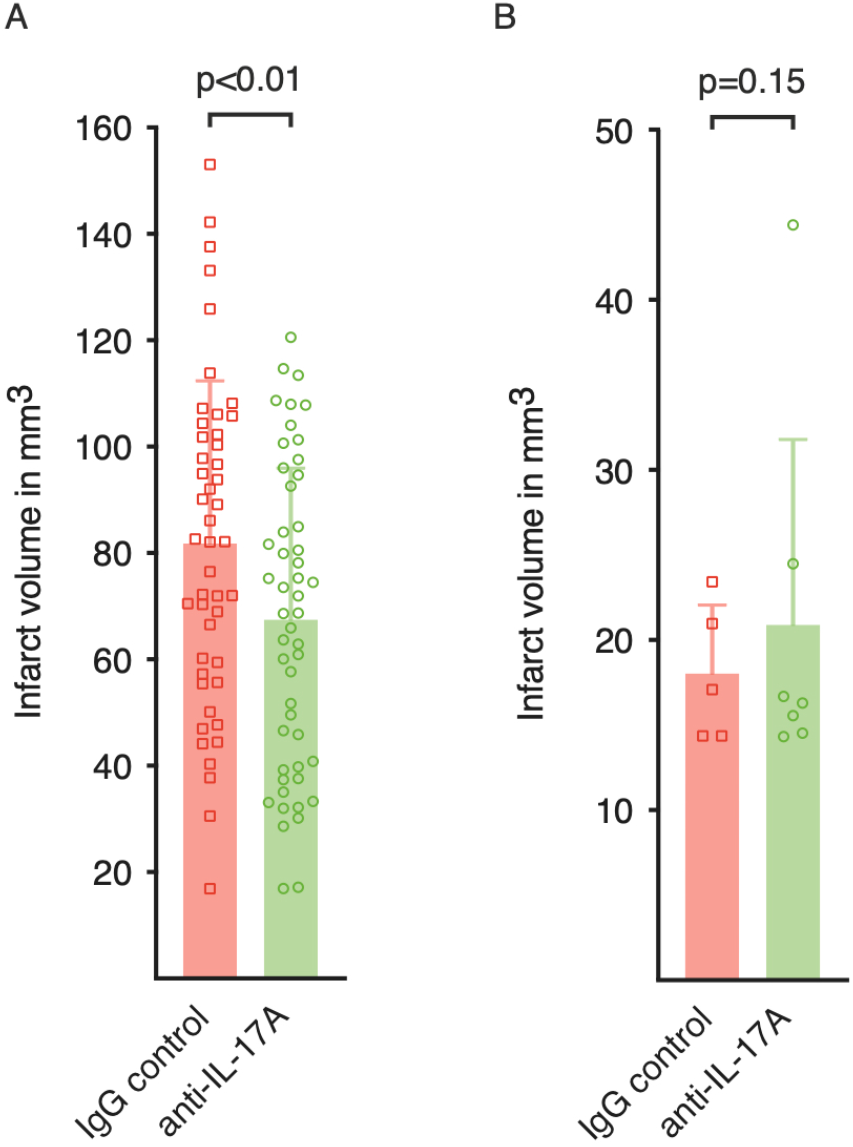
IL-17A neutralization reduced only cortical infarcts. (A) Analysis of combined cortical and striatal infarct volumes (pooled data set). Infarct volumes of rater one are shown in mm3. Graphs show mean±SDs. Statistical analysis was assessed by linear-mixed model analysis (F_(1,185)_=8.91, p<0.01, n=97). (B) Analysis of purely striatal infarct volumes (F_(1,16)_=2.26, p=0.15, n=12)).

The activation of the IL-17A axis can be tracked by the analysis of neutrophil infiltration(Gelderblom et al., 2012). We assessed infiltrating neutrophils in different brain regions on day three following tMCAO by flow cytometry. The analysis revealed a significant relative (Figure 4A) and absolute (Figure 4B) increase of neutrophils in cortical areas compared to striatal areas. We also investigated neutrophil numbers in postmortem brain tissue of mice from our multicenter trial. Brains were sectioned, and absolute neutrophil numbers were quantified following immunohistochemical Ly6G staining. Again, we found in the IgG control group that neutrophils predominantly migrated into cortical areas. When we compared the anti-IL17A with the IgG control group, we found a highly significant inhibitory effect of the anti-IL-17A treatment on the neutrophil infiltration in the cortex but not the striatum (Figure 4C, D). In order to demonstrate the translational relevance of our findings, we examined neuropathological samples from a patient who had died 24 h after having a stroke. In this tissue, H&E staining showed cells with unequivocal physical characteristics of neutrophils (polymorphonuclear) and few monocytes in the edematous parenchyma (Figure 4E). In summary, we showed that the anti-IL-17A treatment is predominantly effective in cortical infarcts and that it results in reduced migration of neutrophils into cortical areas.

**Figure 4.**
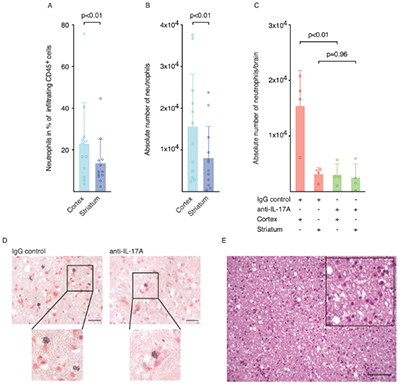
IL-17A neutralization diminished neutrophil infiltration into the cortex. (A) Percentages of neutrophils in infiltrating CD45^+^ cells and (B) absolute numbers of neutrophils in cortical and striatal brain tissue three days following tMCAO. Graphs show mean±SDs. (Students t-test for dependent variables, (A) t_(11)_=3.68, p<0.01, n=12; (B) t_(11_)=3.8, p<0.01, n=12). Cell counts were determined by flow cytometric analysis of CNS-infiltrating cells. (C) Quantitative analysis of Ly6G^+^ cell counts per ipsilateral hemisphere in representative animals of our trial. Graphs show mean±SDs. Statistical analysis was performed with a two-factorial analysis of variances with repeated measures and Šidák’s multiple comparison tests for pairwise comparisons (Cortex: t_(12)_=4.85, p<0.01, Striatum: t_(12)_=0.24, p=0.965, n=8)). (D) Immunohistochemical staining of neutrophils (Ly6G) in mice from both treatment groups 3 days after tMCAO (scale bar 20 µm). (E) H&E staining of ischemic human brain tissue 24 hours after stroke (scale bar 100 µm).

### The gut microbiome differs in between the four centers

The gut microbiome influences the immune response and the overall outcome following stroke (Xia et al., 2019). In order to determine differences in the composition of the microbiome and thus a possible influence on stroke outcome, we sequenced the variable regions v3 and v4 of the 16S rRNA of the gut bacteria. Specimens of 10 specific pathogen free (SPF) mice of each center were collected. The analysis of the operational taxonomic units (OTUs), revealed significant differences in the species diversity between most study centers (Figure 5A, B). Furthermore, the principal coordinate analysis showed distinct clusters, clearly distinguishing the four test centers (Figure 5C).

**Figure 5.**
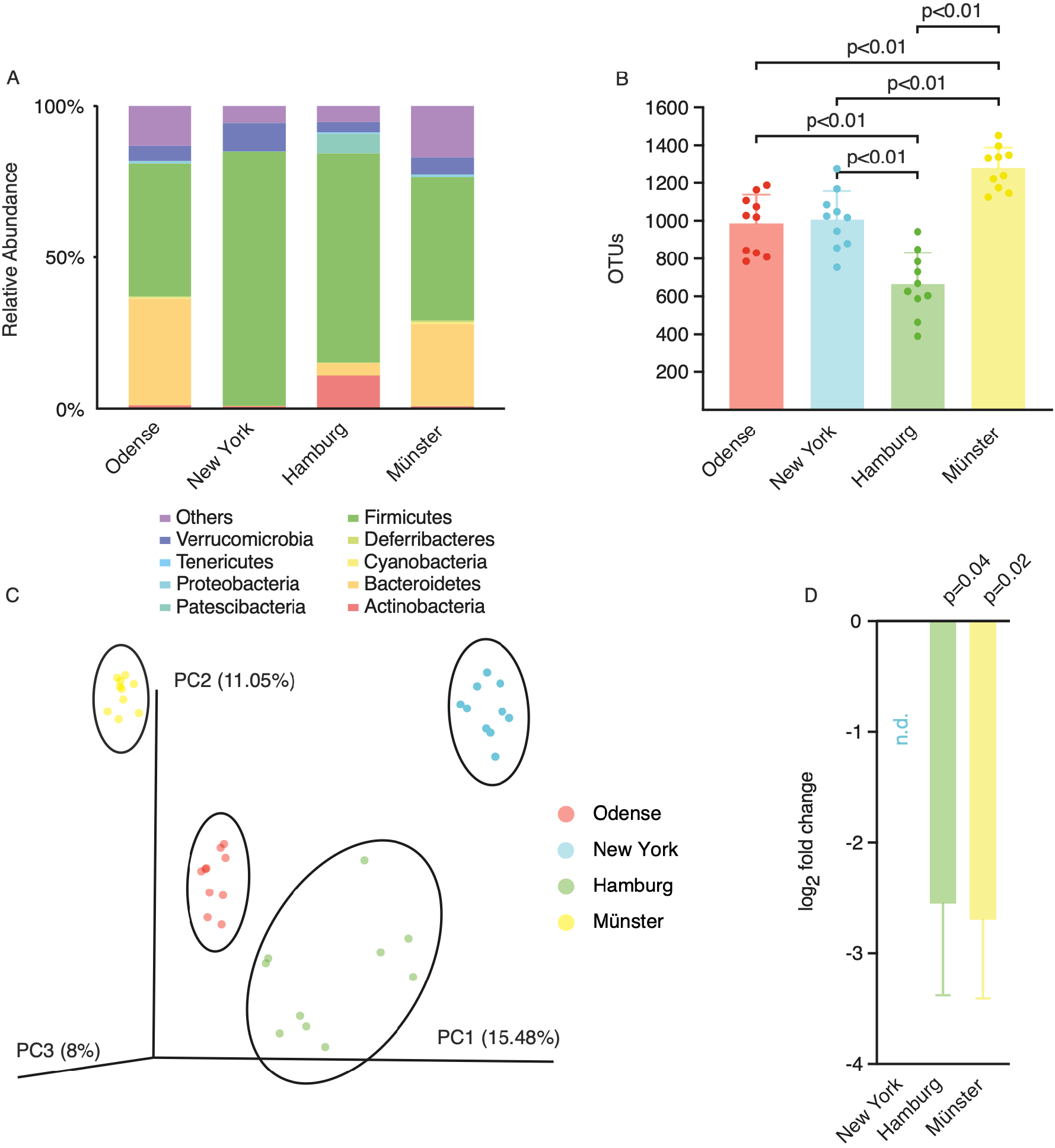
The microbiome differs between the four centers. (A) Taxonomic summary of OTUs on phylum level, n=10 in each center. (B) Total numbers of OTUs for each center. Graphs show mean±SDs. Statistical significance was assessed by pairwise t-tests for independent samples (New York vs. Odense: t=0.29, p=1; New York vs. Hamburg, t=4.76, p=0.001; New York vs. Münster: t=-4.55, p=0.004; Hamburg vs. Odense: t=-4.48, p=0.002; Hamburg vs. Münster: t=-9.65, p<0.001; Münster vs. Odense: t=-4.88, p=0.001, n=10 in each center). (C) Principle coordinate analysis (PCoA) of the family composition. Clusters of each center are highlighted with an ellipse. Statistical testing was performed with an analysis of similarities (R=0.78, p<0.001, n=40). (D) qPCR analysis for SFB presence in mice from the four different centers. Expression levels were normalized to corresponding SFB levels in mice from Odense. Shown is the log2 fold change as means±SDs. SFB were not detectable in the New York cohort. Statistical significance was assessed by t-tests for independent samples (Hamburg: t_(14)_=2.23, p=0.04, n=16; Münster: t_(16)_=2.58; p=0.02, n=18).

The single center analyzes of infarct sizes had revealed that one center (Odense) showed no effect of the IL-17A treatment (Supplemental Figure 3A, B). The colonization with segmented filamentous bacteria (SFB) can induce IL-17A producing T cells (Ivanov et al., 2009). SFB were assessed by quantitative real-time PCR in the fecal material. The qPCR showed that animals from Odense had significantly higher amounts of SFB compared to Münster and Hamburg, while in New York no SFB were detectable (Figure 5D). In summary, the data showed that the microbiome and the SFB levels significantly differed between the four test centers and that the colonization with SFB correlated with a worse response to IL-17A neutralization.

## Discussion

This pRCT provides evidence that IL-17A neutralization results in smaller infarcts and improved survival. These findings confirm the results from single center studies investigating the impact of IL-17A on stroke outcome (Benakis et al., 2016; Gelderblom et al., 2012; Shichita et al., 2009). Recent consensus statements have indicated the need for pRCTs to validate and eventually translate experimental research. Our trial followed the given guidelines (Dirnagl et al., 2013) and is the first positive pRCT in stroke using the tMCAO model.

There are several experimental stroke models available: the tMCAO, the permanent MCAO (pMCAO), and the photothrombosis (ptMCAO) model (Sommer, 2017). tMCAO resembles the occlusion/reperfusion injury of a proximal middle cerebral artery (MCA) infarct in humans while pMCAO and ptMCAO are similar to distal cortical infarcts of the MCA. The advantage of pMCAO and ptMCAO is a smaller variability in stroke size and, therefore, smaller effects can be detected more easily (Lipsanen & Jolkkonen, 2011). Compared to the human situation, the downsides of these two models are the artificial opening of the dura mater (pMCAO) or the use of the radical inducing chemical rose Bengal (ptMCAO). tMCAO, as in our trial, has a much higher variability including cortical and subcortical infarcts but corresponds better to the variability seen in human trials (Elkins et al., 2017; Llovera et al., 2015; Lovblad et al., 1997). This dilemma is also documented in the only other pRCT in experimental stroke research. In this trial, Llovera and colleagues investigated the effects of CD49d-specific antibodies (Llovera et al., 2015). A significant reduction of infarct volume was only documented for pMCAO but not for tMCAO. Subsequent human trials (ACTION 1 and 2) using CD49d-specific antibodies failed to find any effect on stroke progression (Elkind et al., 2020; Elkins et al., 2017). However, the range in infarct volume with 1 - 212 ml in the clinical study compared better to the non-significant tMCAO (0.001 - 0.095 ml) but not to the significant pMCAO results (0,003-0,016 ml) (Elkins et al., 2017; Llovera et al., 2015). Consequently, two different approaches seem feasible. First, only therapeutics, which result in significant beneficial effects despite a high variability in a pRCTs, should enter human stroke trials. Second, should a therapeutic have a significant benefit in a pRCT with low variability (e.g. pMCAO, ptMCAO), the corresponding human stroke trial should use inclusion criteria ensuring low variability. In our anti-IL-17A trial, the range in stroke volume was 0.0013 - 0.15 ml, which corresponds to the variability of stroke volumes in most human studies (Elkins et al., 2017; Ganesan, Ng, Chong, Kirkham, & Connelly, 1999; Lovblad et al., 1997).

The main physiological function of IL-17A is to induce innate immune responses towards pathogens resulting in a rapid upregulation of neutrophil attracting chemokines (reviewed by (McGeachy, Cua, & Gaffen, 2019; Veldhoen, 2017)). In our study, we could demonstrate that the neutralization of IL-17A led to a diminished migration of neutrophils to the brain. Interestingly, the overall infiltration of neutrophils and their inhibition by anti-IL-17A treatment was much more prominent in ischemic cortical areas than subcortical/striatal areas. This finding could explain why IL-17A neutralization was only effective in animals with cortical infarcts. A potential reason for this tropism of neutrophils to cortical areas, is that IL-17A^+^ γδ T cells are mostly located in the meninges (Alves de Lima et al., 2020; Benakis et al., 2016; Ribeiro et al., 2019).

However, IL-17A signaling in the brain does not only have detrimental effects. IL-17A from meningeal γδ T cells can support short-term memory and synaptic plasticity through induction of glial brain-derived-neurotrophic factor (BDNF) (Ribeiro et al., 2019). This dichotomy of IL-17A effects in the CNS is likely explained by the micro milieu in affected target tissues. Net outcomes of the IL-17 receptor activation strongly depend on the cellular target and the cooperative stimulation by other cytokines. This cooperation results in synergistic activation of downstream signaling pathways (Ruddy et al., 2004). In particular, IL-17A synergizes with strong NF-κB activators, such as IL-1β and TNF-α, which are both rapidly upregulated in ischemic brain tissue by microglia and macrophages (Clausen et al., 2016; Yli-Karjanmaa et al., 2019). The synergistic stimulation of IL-17 receptor positive target cells, such as astrocytes, by IL-17A and IL-1β and/or TNF-α induces an amplification of pro-inflammatory factors in the ischemic brain (Gelderblom et al., 2012). Once the acute injury has subsided, the decrease in synergistic pro-inflammatory cytokines might result in more neuro-regenerative effects of IL-17A (Lin et al., 2016). For this reason, a therapeutic interference with anti-IL17A antibodies should be of short duration and limited to the acute stage of stroke.

In the ischemic brain of the mouse, we could show that IL-17A is produced by vγ6^+^ γδ T cells (Arunachalam et al., 2017). These data and our current study have the drawback that they stem from studies in young male mice, which do not reflect the human stroke population. Human stroke patients are older and typically suffer from comorbidities such as atherosclerosis, hypertension, and diabetes. In addition, the cellular source of IL-17A in humans is less well understood. In human autoimmunity, IL-17A producing immune cells encompass CD8^+^ T cells, Th17 cells, NK cells, NKT cell, neutrophils, and mast cells (reviewed by (Lubberts, 2015)). However, the IL-17A pathway is highly conserved between mammals and downstream pathways such as chemotaxis are comparable (Veldhoen, 2017). We could show that IL-17A producing cells and neutrophils infiltrate ischemic human brains at early stages (Gelderblom et al., 2012). These facts provide clear evidence that the IL-17A axis contributes to human stroke.

Another strong modulator of immune responses is the microbiome (Kamada, Seo, Chen, & Nunez, 2013). The microbiome has the potential to exert both pro- and anti-inflammatory responses and is intimately linked to the proper function of our immune system (Kamada et al., 2013; Round & Mazmanian, 2009). This influence is also observed on γδ T cells. The microbiome impacts their proliferation, activation as well as their IL-17A secretion (Singh et al., 2016; Zhang et al., 2019). Particularly, segmented filamentous bacteria (SFB) shape the IL-17A polarization. Accordingly Ivanov et al. could show that there is a shift towards increased Th17 cells in the gut of mice colonized by SFB (Ivanov et al., 2009). In stroke, recolonizing germ-free mice with dysbiotic microbiota increases lesion volume through a proinflammatory T cell polarization (Singh et al., 2016). Even further, Benakis et al. demonstrated that the gut microbiome influences the presence of IL-17A and the outcome following stroke (Benakis et al., 2016). Taken together, there is clear evidence that the microbiome has a substantial impact on the IL-17A response and should be considered in any study dealing with IL-17A neutralization or even any immune modulation. In our study, we can demonstrate substantial heterogeneity within the microbiome among the four participating centers. This reflects the heterogeneity in human cohorts (Hall, Tolonen, & Xavier, 2017) and does not eliminate the effect of anti-IL-17A treatment. However, when we analyzed the presence of SFB, we found that the center with the highest SFB level showed no effect of the IL-17A neutralization. This finding suggests that higher levels of IL-17A might diminish the effectiveness of our IL-17A antibody, at least of the applied dosage. Thus, an adaptation of the anti-IL-17A dosage might be required in individuals with constitutively elevated IL-17A levels.

The last 20 years have seen two milestones in stroke treatment: i.v. thrombolysis (National Institute of Neurological & Stroke rt, 1995) and more recently the mechanical recanalization (Berkhemer et al., 2015). While the reconstitution of the blood flow has certainly been helpful in a subgroup of stroke patients, still only a minority benefits from the treatment. Therefore, it is crucial that new targets are being developed. We believe that IL-17A could be such a promising target. Our pRCT trial offers the best possible preclinical data basis to design a trial in humans with strokes due to large vessel occlusions. In this scenario anti-IL-17A treatment could follow endovascular thrombectomy.

## Material and methods

### Study design

The study was performed between 2017 (initiation) and 2018 (unblinding) in an international consortium consisting of four test centers. The infarct size was the primary endpoint. Secondary endpoints were mortality, neurological outcome and neutrophil infiltration. The Hamburg study center (M.G.) initiated the trial, coordinated the study design and performed central data analysis. Hamburg (M.G.), New York (J.A.), Odense (B.C.) and Münster (J.M.) performed the tMCAO experiments. Sample size per stroke model was determined a priori by performing a power analysis (Power=0.95, G-Power). The group sizes were based on published data.

### Mice

C57Bl/6 mice were provided by different suppliers (Münster and Hamburg: Charles River, Odense: Taconic, New York: The Jackson Lab) and kept in the respective animal facility at least two weeks before experiments started. We used male mice with an age of 11 to 13 weeks and a bodyweight of 24 to 28 g. In total, 136 mice were randomized and treated. The animals were housed under the local guidelines of each study center and assessed at least once a day.

### Transient middle cerebral artery occlusion (tMCAO)

Temporary middle cerebral artery occlusion was achieved by using the intraluminal filament method (Doccol: Filament # 602112PK10) as previously described for 45 min (Gelderblom et al., 2009). All mice were monitored for heart rate, rectal body temperature, and cerebral blood flow with the use of transcranial temporal laser Doppler scans. The cerebral blood flow in the area of the MCA showed a reduction by 80%, which did not differ between groups (data not shown). Heart rate, and rectal body temperature also showed no difference in our cohorts. After stroke induction, every mouse was repeatedly scored on a scale from 0 to 5 (0, no deficit; 1, preferential turning; 2, circling; 3, longitudinal rolling; 4, no movement; and 5, death) immediately after reawakening and every day until killing. Mice were sacrificed three days after tMCAO and perfused with phosphate-buffered saline (PBS) and paraformaldehyde (PFA) 4% afterward. Mice that did not survive until day three were excluded for the further analysis of the primary outcome parameter. Brains were kept in 4% PFA for 24h, stored in PBS, and sent to Hamburg for further analysis.

### Infarct volume assessment by Magnetic Resonance Imaging

The measurements of the specimens from Odense, New York and Münster were performed on a 7 Tesla system (ClinScan, Bruker). The specimens from Hamburg were measured on a 9,4 Tesla system (BioSpec, Bruker). For the MRI acquisition, brains were placed in 15 ml PBS filled falcons. The imaging protocol comprised an apparent diffusion coefficient (ADC) and diffusion-weighted images (DWI). Calculations of infarct volumes were performed on ADC images using ImageJ (NIH). Infarct sizes were corrected for brain edema (Gerriets et al., 2004). The rating of the infarct volumes was performed by two independent and blinded raters. The comparison of the infarct sizes of the two investigators revealed a highly significant correlation (Supplemental Figure 4). MRI scans and all analysis files were stored on a central database.

### Immunohistochemistry

Following MRI, the PFA fixed brains were embedded in paraffin. Coronal sections of 10 µm thickness were collected every 400 μm. For the immunostaining of neutrophils, we performed antigen retrieval in 10 mM citrate buffer (pH 6.0) and blocked with normal rabbit serum. Ly-6G clone 1A8 (1/500, Biolegend) was incubated overnight at 4°C. Vectastain Elite ABC HRP kit (Rabbit-anti-rat secondary AB, 1/200) was used for visualization of the secondary antibody. For nuclear staining, Nuclear Fast Red (SigmaAldrich) was added for 5 minutes on RT. Images were acquired on a digital slide scanner (Hamamatsu; Nano Zoomer). Manual cell counting was done with the NDP viewer (Hamamatsu). Individual neutrophils on the entire slide were marked by hand in order to avoid double counts. Assignment to either the cortical or striatal group was assessed by colocalization to the respective anatomical structure. Total numbers of neutrophils were calculated according to the following calculation: Total number of neutrophils = N(n_1_+n_2_+…n_i_) x 40 (N = Sum of all counted neutrophils in the brain, n_i_ = Number of neutrophils counted in each slide_i_ and 40 = Number of slides of 10 µm thickness in the position 1…i). For each brain, neutrophils were analyzed in 15 slides. Images of representative neutrophils in both groups were taken with a transmission light microscope (Apotome, Zeiss) and a 63x plan-apochromatic oil DIC objective. For analysis of autoptic human brain tissue, cases were selected from the files of the Institute of Neuropathology at the University Medical Center Hamburg-Eppendorf. Brain specimens had been fixed in 4% buffered formalin for at least two weeks before paraffin-embedding. Brain sections (3 m thick) were stained according to standard haematoxylin and eosin stain procedures.

### Collection of feces

We collected feces of 10 mice in each center for the analysis of the gut microbiome. All mice housed for at least two weeks in the animal facility of the respective centers in order to adapt to local microbiota. The specimens were collected early in the morning directly from each mouse in order to avoid the loss of anaerobic bacteria. Immediately after collection, the feces were homogenized in 96% ethanol to stop DNA degradation.

### Nucleic Acid Extraction

Nucleic acid (DNA and RNA) was extracted from 200 µl sample volume using an automated extraction system, Qiasymphony (Qiagen, Hilden), according to the manufacturer’s instructions.

### 16S rRNA amplicon library preparation, MiSeq sequencing, and bioinformatic analysis

V3–V4 region 16S rRNA amplicon sequencing was performed as recently published (Lamprecht et al., 2019). The following primers containing the Illumina adapter consensus sequence were used: F: 5’-TCGTCGGCAGCGTCAGATGTGTATAAGAGACAGCCTACGGGNGGCWGCAG-3 and R: 5-GTCTCGTGGGCTCGGAGATGTGTATAAGAGACAGGACTACHVGGGTATCTAATCC-3. Samples were multiplexed using the Illumina Nextera XT Index Kit and sequenced by 500 PE sequencing on the MiSeq platform. FastQC (Babraham Bioinformatics, Babraham Institute, UK) was used to determine the average quality scores of each sample before and after paired reads. The paired ends in each sample were joined, and all sequences less than 250 bp and/or with a Phred score <33 were discarded. Quality filtering was applied using QIIME 53 (at Phred ≥ Q20). We performed operational taxonomic unit (OTU) clustering and alpha- and beta-diversity analysis using QIIME 2 (Bolyen et al., 2019). A chimera filter was applied using USEARCH 8.1. All sequences were clustered based on 97% similarity to reference sequences. The reads that did not meet the 97% similarity criteria were clustered de novo. Taxonomy levels of representative sequences in the OTU table were assigned at 95% similarity based on the SILVA database. We calculated alpha diversity based on the total amount of OTUs. Analysis of beta diversity statistics (analysis of similarities, ANOSIM) was performed to determine if differences between the distributions of microbiota profiles were significant.

### Quantitative PCR analysis of Segmented Filamentous Bacteria (SFB)

RT-PCR was performed on 7 ng of purified fecal DNA in three technical replicates. PCR was performed with Power SYBR Green PCR Master Mix (Thermofischer) on a Lightcycler 96 (Roche) with the following primers: SFB 736F: GACGCTGAGGCATGAGAGCAT, SFB 844R: GACGGCACGRATTGTTATTCA, and universal bacterial r16S gene primers 16S-V2-101F: AGYGGCGIACGGGTGAGTAA and 16S-V2-361R: CYIACTGCTGCCTCCCGTAG. Ct values of SFB amplicons were normalized to r16S gene Ct values by the ΔΔCt method and data are expressed as fold change over SFB levels in mice from Odense as reference group. Analysis was performed with Excel (Microsoft) and GraphPad Prism.

### Flow Cytometric analyzes

Fluorescence-activated cell sorting (FACS) measurements were performed with an LSR Fortessa (BD Biosciences). Data analysis was done with FlowJo (BD Biosciences). After PBS perfusion, the removed brains were cut into 1 mm thick slices, and the cortex was dissected from the striatum under a microscope. Only ipsilesional hemispheres were taken, digested for 30 min at 37°C (1 mg/ml collagenase, 0.1 mg/ml DNAse I in DMEM), and pressed through a 40 µm cell strainer. Cells were incubated with standard erythrocyte lysis buffer on ice and separated from myelin and debris by Percoll gradient (GE Healthcare) centrifugation. All antibodies were used with a dilution of 1/100. Incubation was performed for 30 minutes at 4°C. For cell counting, BD Trucount (BD Biosciences) tubes were taken. The following antibodies were used: Biolegend: TCR γ/δ (Clone GL3), CD11b (M1/70), CD3 (17A2), B220 (RA3-6B2), F4/80 (BM8), CD8 (53-6.7), NK1.1 (PK136), CD4 (RM4-59), eBioscience: MHCII (M5/114.15.2), CD11c (N418), CD45 (30-F11), BD Bioscience: Ly6G (1A8).

### Statistical Analysis

We computed a generalized linear mixed model analysis (SPSS, IBM, © 2019) to test for treatment effects on the infarct volume. This model allowed us to account for two raters as random effects. Fixed effects were set as treatment and center in order to compute pairwise contrasts between the treatment groups for each center. Normal distribution was checked beforehand by histogram analysis. For the pairwise comparisons, p-values were adjusted with Šidák’s correction in order to account for multiple testing. Each specimen was divided into one of the two subgroups (with- and without cortical infarction) by two independent raters. Within each subgroup, we conducted the same analysis as we did in the main group. Pearson’s r was calculated for interrater reliability. Mortality was analyzed with a log-rank test. Neuroscores were analyzed with a Mann-Whitney test for non-parametric data and bodyweight was analyzed using a mixed model analysis to account for missing values with time points and treatments as fixed effects. Šidák’s test was assessed to test for pairwise contrasts of the two treatment groups on each time point. Flow cytometric data on neutrophil infiltration was tested using a two-sided student’s t-test for dependent samples. Neutrophil counts from immunohistochemical studies were analyzed with a multivariate analysis of variances with repeated measurements of variances with compartments and treatments as independent variables. Šidák’s test was assessed to check for pairwise contrasts of the two treatments in the two compartments. Differences in the OTUs were assessed with t-tests for independent samples (QIIME 2). An analysis of similarities was performed to test for significance in the beta-diversity (QIIME 2). RT-qPCR data were analyzed with student’s t-tests for independent samples. In all tests, the alpha level was set on 0.05. If not stated otherwise GraphPad Prism 8 was used for the analysis.

## Appendix

## Study approval

We conducted all experiments according to the Guide for the Care and Use of Laboratory Animals published by the US National Institutes of Health (publication No. 83–123, revised 1996) and performed all procedures in accordance with the ARRIVE guidelines (Animal Research: Reporting of In Vivo Experiments; http://www.nc3rs.org/ARRIVE). All animal experiments were approved by the respective local animal care committees.

## Author contributions

M.G. initiated this trial; M.G., T.M., J.M., J.A., B.H.C., T.V.A., G.T., and C.G. designed the study protocol, supervised the study, reviewed, and verified original data sets. S.K. performed the central analysis of infarct sizes. S.K., J.K.S., C.J., L.G.B., P.L., C.B., C.F., J.S. performed experiments. S.K. and H.P. performed central statistical analysis. M.G., S.K., T.M., J.M., B.H.C., J.A., G.T., and C.G. wrote the manuscript.

## Acknowledgments

We thank Oliver Schnapauff and Lennart Pöls for excellent technical assistance, Karoline Degenhardt for the immunohistochemical Ly6G staining, Nicole Fischer for the analysis of the microbiom data, Christoph Debus for the analysis of infarct sizes and the FACS Core Unit for the technical support. This work was supported by grants of: Deutsche Forschungsgemeinschaft (KFO Immunostroke 2879: Project A1 (TM 4375/5-1), Project A3 (TM 4375/6-1), Project B3 (MG 2036/1-1)); Deutsche Forschungsgemeinschaft SFB 1382 (TM Project A13); The Schilling foundation (TM); The core unit PIX of the Interdisciplinary Center for Clinical Research Münster.

## Supplemental Material

Supplemental figure 1:

Study design, attached separately

**Supplemental Figure 2.**
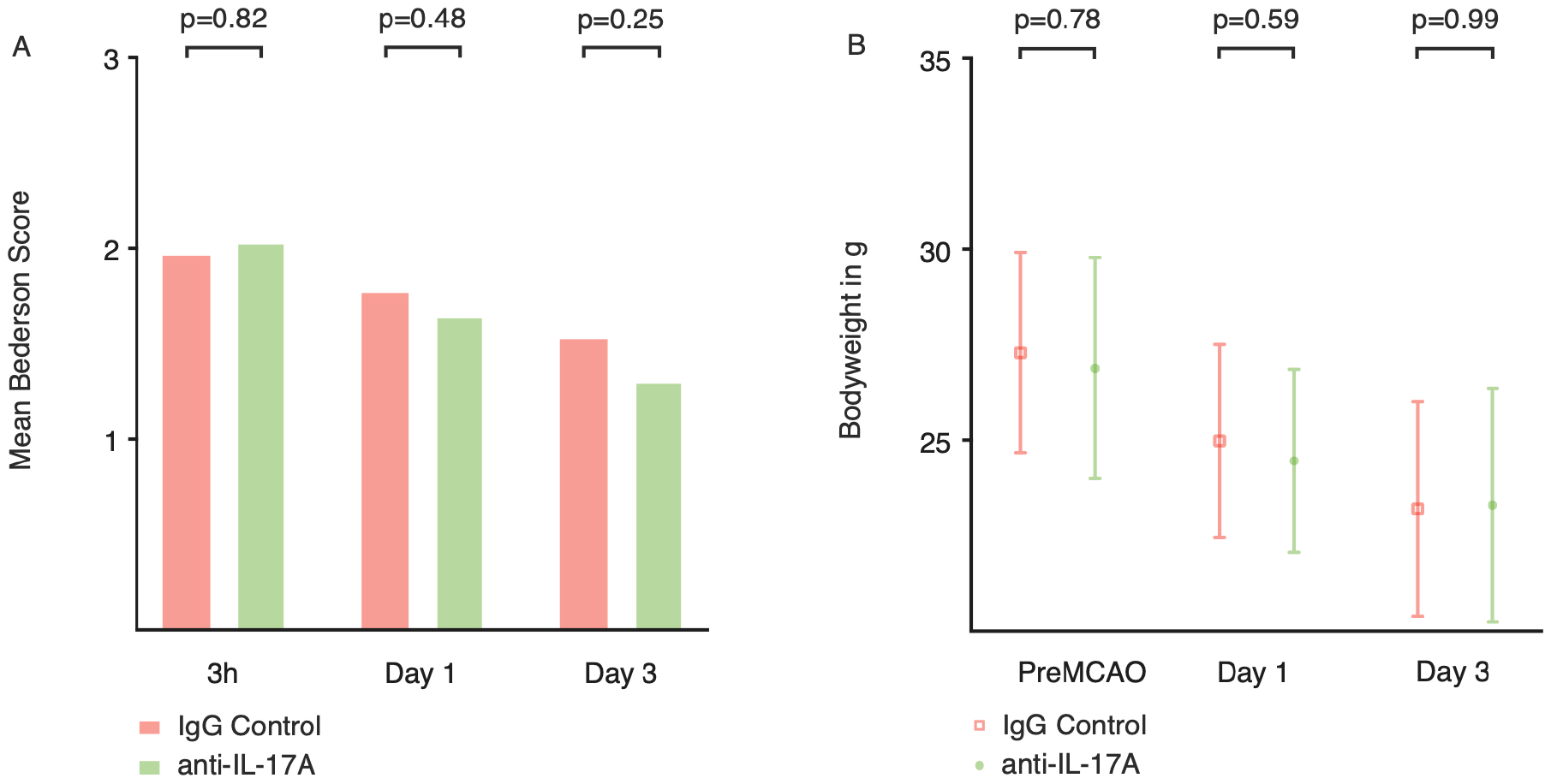
Bederson Neuroscore and bodyweight as secondary outcome parameters. (A) Neurological scores at 3h and at days 1 and 3 of animals that were treated with 500 µg isotype control or 500µg of anti-IL-17A. Significance was assessed by Mann-Whitney-U Test (3 hours: U=1878, p=0.82, n=62 in both groups, Day 1: U=1417, p=0.48, n= 59 in the anti IL-17A group and n=52 in the IgG control group. Day 3: U=1341, p=0.25, n=60 in the anti IL-17A group and n=51 in the IgG control group). (B) Bodyweight in g of mice of both treatment groups. Statistical analysis was performed with a linear mixed model analysis and Šidák’s multiple comparison tests for pairwise comparisons (PreMCAO: t_(125.4)_=0.84, p=0.78, n=128. Day 1: t_(109.4)_=1.14, p=0.59, n=114. Day 3: t_(106.3)_=0.17, p=0.99, n=109).

**Supplemental Figure 3.**
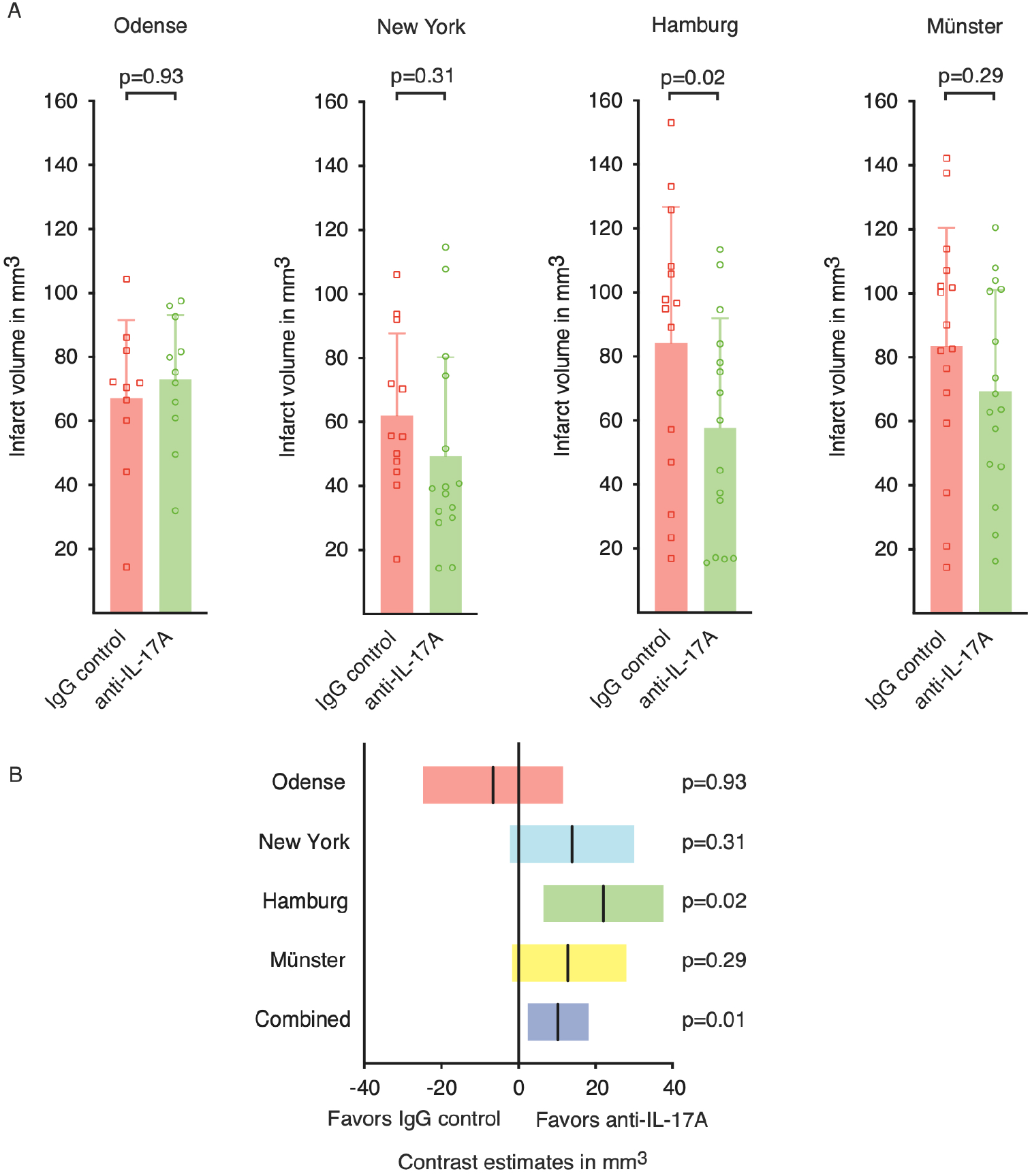
Infarct size divided into individual centers. (A) The infarct sizes of rater one in mm^3^ for each individual center. Infarct volumes were analyzed three days following tMCAO by MRI. Graphs show mean±SDs. Statistical significance was assessed by linear-mixed model analysis with pairwise comparisons. (B) Pairwise comparisons of infarct volumes for single centers. Post-hoc Šidák adjustments were made for the alpha level. The estimated difference is shown in mm^3^±95% CI (Odense: F_(1,209)_=0.51, p=0.93, n=21; New York: F_(1,209)_=3.04, p=0.31, n=27; Hamburg: F_(1,209)_=7.76, p=0.02, n=29; Münster: F_(1,209)_=2.88, p=0.29, n=32).

**Supplemental Figure 4.**
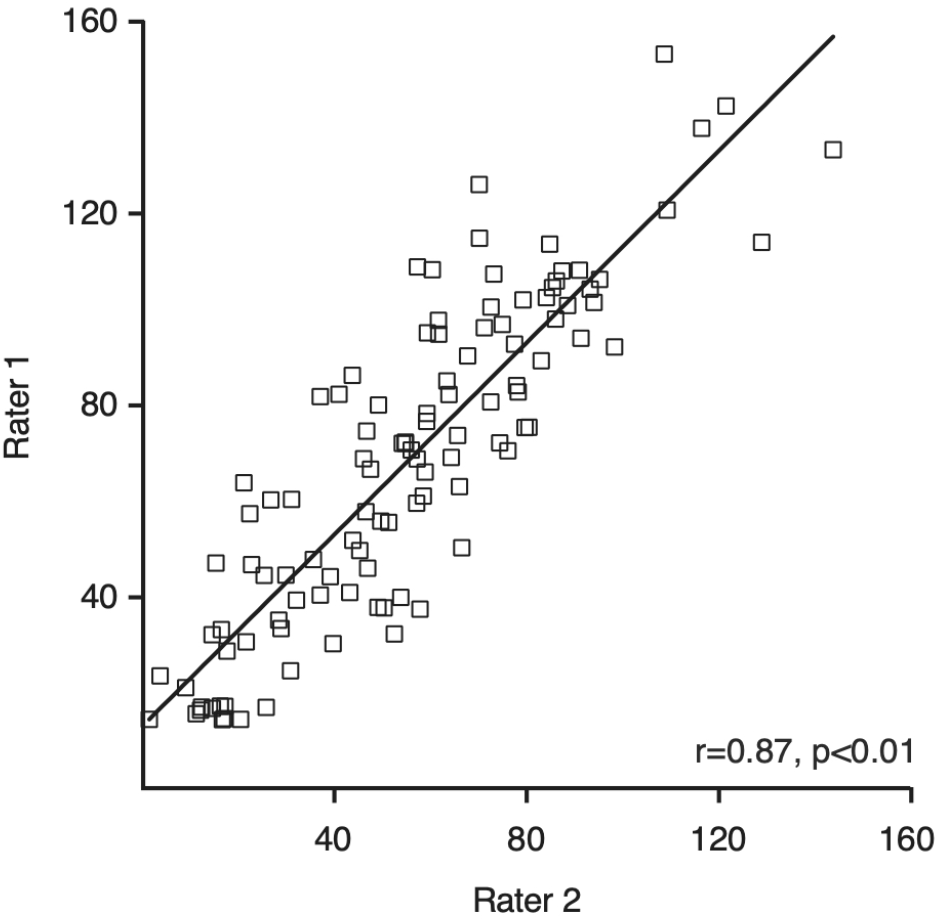
Comparison of the volumetry of the raters. Correlation of MRI based ratings of infarct volumes of rater one and two (Pearson r=0.87, CI = 0.82 – 0.91, p<0.001, n=109 pairs).

