## Supplemental Figure 1 for "A preclinical randomized multicenter trial of anti-IL-17A treatment for acute ischemic stroke"

**Preclinical randomized controlled multicenter trial to assess the efficiency of an anti-IL17A antibody treatment in MCAO**

### 1.1 Background:

Several studies in experimental stroke have shown, that the disruption of the IL-17 axis is protective in stroke[1-3]. In murine models of stroke *Il17a*$^{-/-}$[1] and *Il17ra*$^{-/-}$[2] mice were protected from ischemic tissue damage and anti-IL-17A antibody treatments[2] improved neurological outcomes. Currently available studies were performed in single centers.
To overcome the low reproducibility and translation of pre-clinical single center studies preclinical randomized controlled multicenter trials (pRCT) were proposed. Aim of the current study is to test the efficiency of an anti-IL-17A antibody treatment in a preclinical randomized controlled multicenter stroke trial (pRCT).

Our protocol fulfills fundamental requirements of pRCTs:

- harmonization of experimental protocols among study centers
- a priori sample size calculation
- randomization of the treatment groups
- blinding of all investigators with respect to treatment allocation
- centralized study organization
- analysis by an independent research center

### 2.1. Material provided by Hamburg:

- anti-IL-17A antibody (aliquots a 500µg (3.33 µg/µl), Clone MM17F3; 16.6 mg/kg of bodyweight)

- isotype-control antibody (IgG1) (aliquots a 500µg (3.33 µg/µl), 16.6 mg/kg of bodyweight)

- filament (see below)

### 3. Standardized MCAO procedure:

**see: study design in Figure 2**

### 3.1. Randomization:

Anti-IL-17A antibody and IgG control will be blinded and coded in Hamburg and distributed to each study center.
Antibody treatment, tMCAO, analysis of infarct volume, and secondary outcome parameters will be performed by researchers who are blinded with respect to the treatment groups. Unblinding will be performed after complement of the statistical analyses.

Supplemental figure 1

### 3.2. Power analysis

Power analysis on the basis of previously published effects of an anti-IL-17A treatment in comparison to an isotype ab treatment results in:

t tests - Means: Wilcoxon-Mann-Whitney test (two groups)

| | | |
|---|---|---|
| Analysis: | A priori: Compute required sample size | |
| Input: | Tail(s) | = One |
| | Parent distribution | = Normal |
| | Effect size d | = 1,2 |
| | α err prob | = 0,05 |
| | Power (1-β err prob) | = 0,95 |
| | Allocation ratio N2/N1 | = 1 |
| Output: | Noncentrality parameter δ | = 3,42 |
| | Critical t | = 1,69 |
| | Df | = 30,48 |
| | **Sample size group 1** | **= 17** |
| | **Sample size group 2** | **= 17** |
| | Total sample size | = 34 |
| | Actual power | = 0,9551119 |

### 3.3. Animals:

- C57Bl/6 mice
- males
- age 11-13 weeks
- weight 24-28 g
- strokes should be performed between 8.00 a.m. and 16.00 a.m.

### 3.4. anti-IL-17A or IgG control administration:

- 500 µg / animal
- concentration 500µg in 150 µl
- 1h after reperfusion
- intravenous administration: retro-orbital or tail-vein
- **group size:** **- anti-IL-17A 17 animals**
  **- IgG control 17 animals**

### 3.5. MCAO-procedure:

#### 3.5.1 filament:

- Silicon rubber-coated monofilament for MCAO model.

- Filament size 6-0, diameter 0.09-0.11 mm, length 20 mm; diameter with coating 0.21 +/- 0.02 mm, and coating length 1.5-2 mm. Pack of 10 filaments. Doccol filaments: 6-0- fine MCAO suture L12PK10 (# 602112PK10)

- Filament will be provided by Hamburg

- Filaments can be re-used if not damaged

3.5.2. occlusion of the left MCA

3.5.3. Occlusion time
- 45 min

3.5.4. Temperature control of mice during surgery
- 35.5-36.5°C

3.5.5. Daily measurement of bodyweight

**3.6. Monitoring during MCAO (before removal of the filament)**

3.6.1. If possible - physiological parameters:
- oxygenation
- heart rate
- breathing frequency

3.6.2. in every mouse laser doppler:
- Flow contralateral / Ipsilateral (before removal of the filament)
- only mice are included with a reduction of the laser doppler flow on the ipsilateral side ≥ 80% in comparison to the contralateral side

**4. Neurological Scoring:**

4.1. Bederson Score

- 3h post MCAO
- 1d post MCAO
- 3d post MCAO

**5. Collection of brain on day 3 post MCAO:**

Mice will be scarified on day 3

5.1. - mice will be perfused with
  - 25 ml PBS, 10 ml/ min, 4°C
  - 25 ml PFA, 10 ml/ min, 4°C
  - 24 h post fix in 4 % PFA in PBS, 4°C
  - transfer of brains into PBS, 4°C

5.2. Cooled brains will be sent to HH

**6. Read out parameter**

**6.1. primary read out parameter:**

infarct volumes on day 3 post MCAO will performed by

diffusion weighted MRI (apparent diffusion coefficient (ADC) in Hamburg (**see Fig. 1**)

**6.2. secondary outcome parameter**

6.2.1. mortality

6.2.2. neurological scores

6.2.3. histological analysis of the neutrophil infiltration in ipsilesional hemispheres

### 7 Exclusion criteria for analysis of the primary read out parameter

**7.1 preanalytical exclusion criteria**

7.1.1. Death before reaching the primary endpoint

7.1.2. Body temperature outside the accepted range

7.1.3. reduction of the laser doppler flow on the ipsilateral side < 80% in comparison to the contralateral side

**7.2 analytical exclusion criteria**

7.2.1 Significant destruction of the brain after removal

7.2.2 Visible parenchymal or subarachnoid hemorrhage

7.2.4. No visible infarct on the MRI

7.2.4 Isolated infarct of the brainstem

Supplemental figure 1

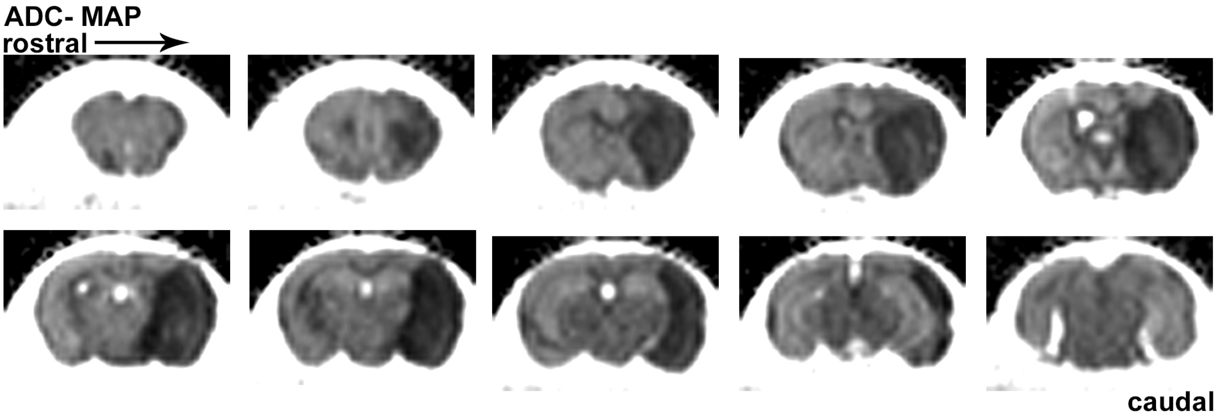


**Figure 1:** Example of infarct size analysis by MRI (DWI) of brains fixed with 4% PFA d3 following tMCAO

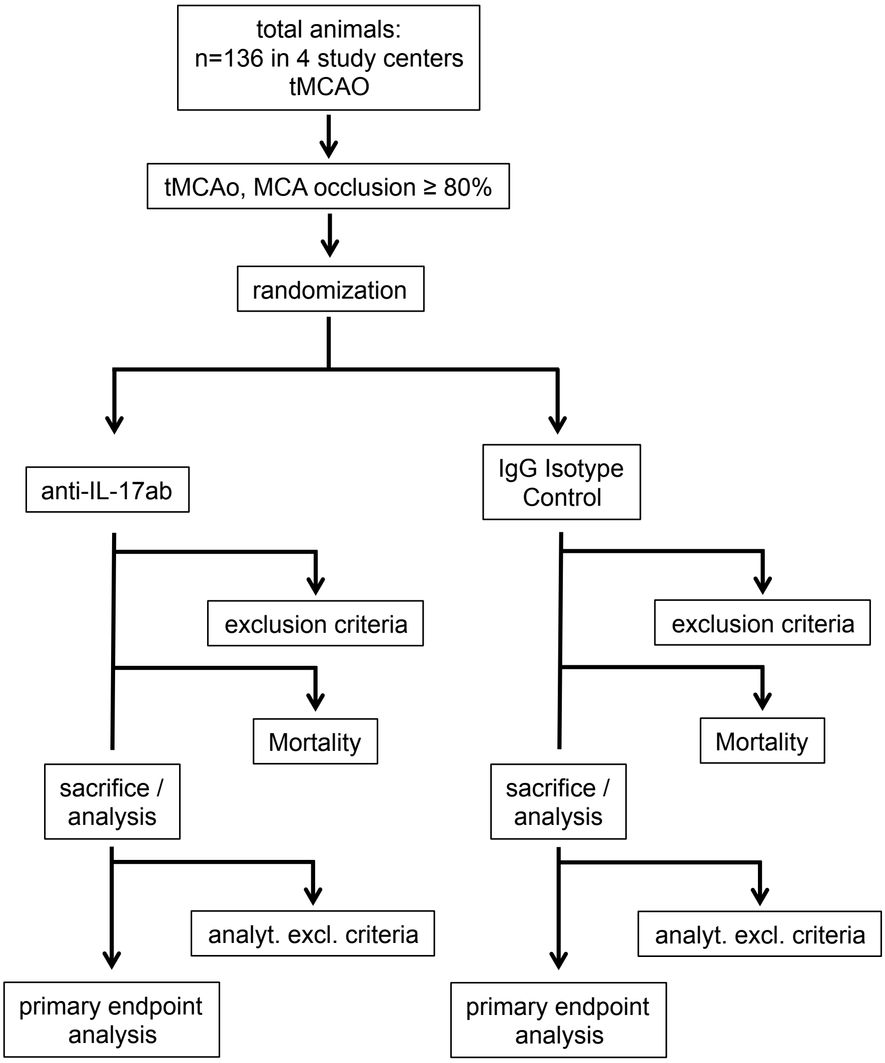


**Figure 2:** Design of the preclinical randomized controlled multicenter trial

Supplemental figure 1
